# Slitflow: a Python framework for single-molecule dynamics and localization analysis

**DOI:** 10.1101/2023.03.01.530718

**Authors:** Yuma Ito, Masanori Hirose, Makio Tokunaga

## Abstract

Single-molecule imaging is a promising method for direct quantification of the dynamics and distribution of biomolecules in living cells. Although numerous methods have been developed to gain biological insights into molecular behavior, the high diversity of microscopes and single-molecule dynamics can result in incomplete reproducibility of analyses. Here, we present Slitflow, an open-source framework for a single-molecule analysis workflow that includes image processing, dynamics analysis, and figure creation. We demonstrated the integrity and flexibility of the workflow using 1) a cherry-picked tracking method combining popular tools and 2) various state-of-the-art analyses in a single pipeline. The software accommodates a large variety of data and methods, paving the way for integrative analyses.

**Code metadata:** 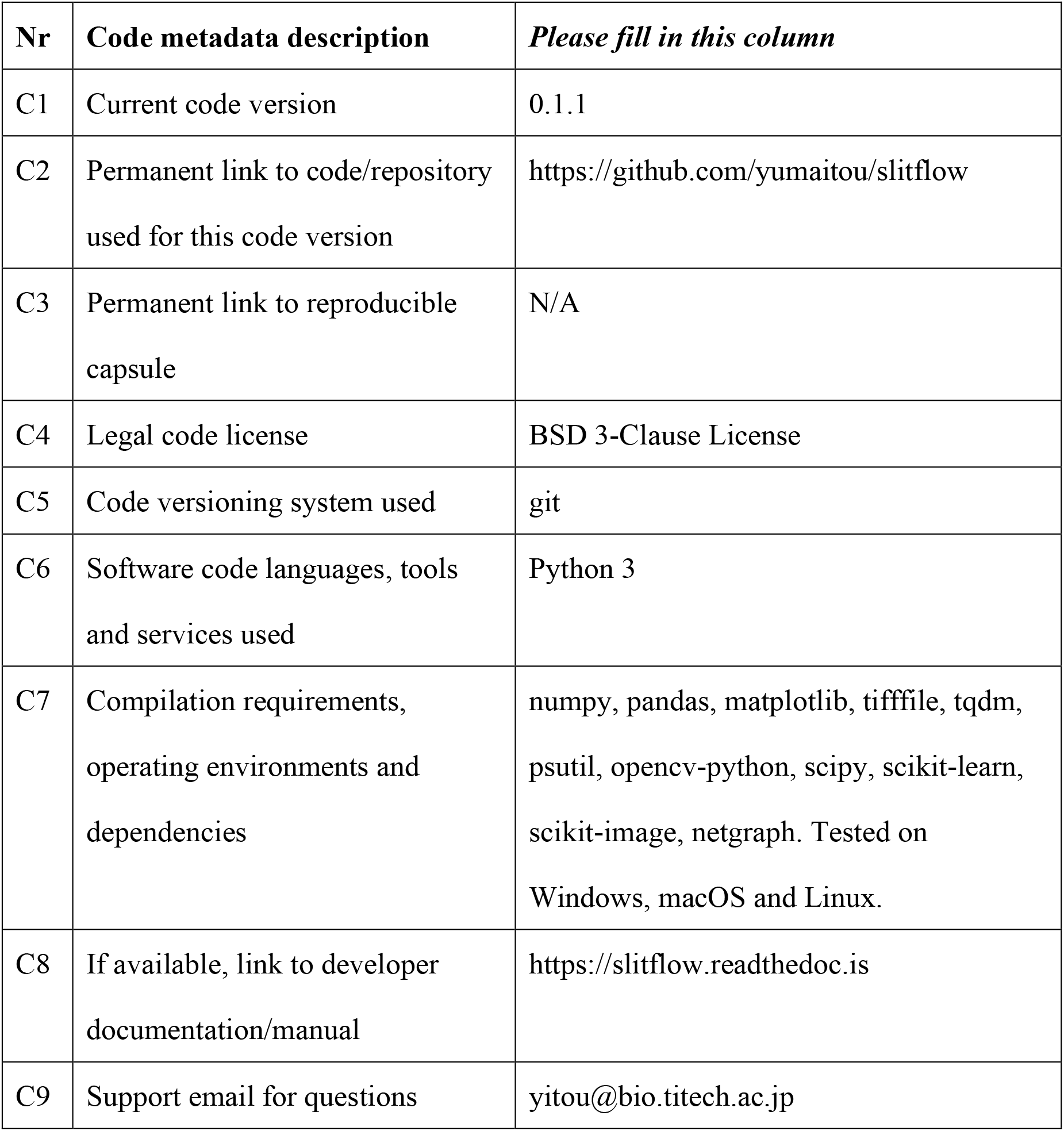

## 1. Motivation and significance

Single-molecule imaging has become a powerful method for understanding the dynamics and distribution of biomolecules in living cells [1, 2], owing to technological advances such as bright fluorophores [3] and specialized illumination [4]. Researchers can obtain various biological insights from heterogeneous single-molecule behaviors using methods developed for trajectory and localization analyses, including physical modeling [5], spatial statistics [6], and machine learning [7]. These methods are provided as script codes, plugins for image-processing software [8], graphical user interfaces [9, 10], and web interfaces [11]. However, users frequently encounter time-consuming problems when converting to software-specific file formats and handling the analysis results, which require programming skills to write additional processing scripts. Additionally, single-molecule images are acquired using highly customized microscopes [1, 2], and molecular dynamics tend to vary widely among targeted protein species [2]. Consequently, optimization of analyses involves not only molecular coordinates but also image processing, detection, and tracking. These steps can complicate the entire analysis workflow and result in an incomplete pipeline, owing to missing parameters and the use of currently unavailable software.

To solve these problems, several software framework packages with graphical user interfaces have been developed [10, 12, 13, 14]. These software packages allow users to construct a robust and reproducible analysis workflow using a well-organized process scheme. However, it is still limited to particular observations, analytical methods, and process steps, thus requiring external programs or intrusive software modifications to add functionality. Consequently, an analytical framework with scalability and integrity is required to create a single pipeline for the whole analysis process of single-molecule imaging. In this paper, we present a comprehensive framework for building a complete single-molecule analysis workflow—namely, Single-molecule Localization-Integrated Trajectory analysis workFLOW (Slitflow). This software was designed to provide reproducibility and accessibility to single-molecule-related processes, from raw data to publication-quality figures. Users can easily create, reproduce, modify, and distribute an integrated workflow using cutting-edge single-molecule analysis methods.

## 2. Software description

### 2.1. Software architecture

Slitflow is a Python package that aims to construct a fully reproducible and universally accessible workflow for single-molecule analysis. To achieve this goal, Slitflow comprises a flexible Data class that executes a task and stores the resulting data. A Data object can be input to the next Data object, the network of Data objects forming the entire workflow of complex single-molecule analysis, from image pre-processing to publication-quality figure creation (**Fig. 1**). This architecture was designed to realize four different levels of reproducibility and accessibility for the three workflow elements described below.

**Fig. 1.**
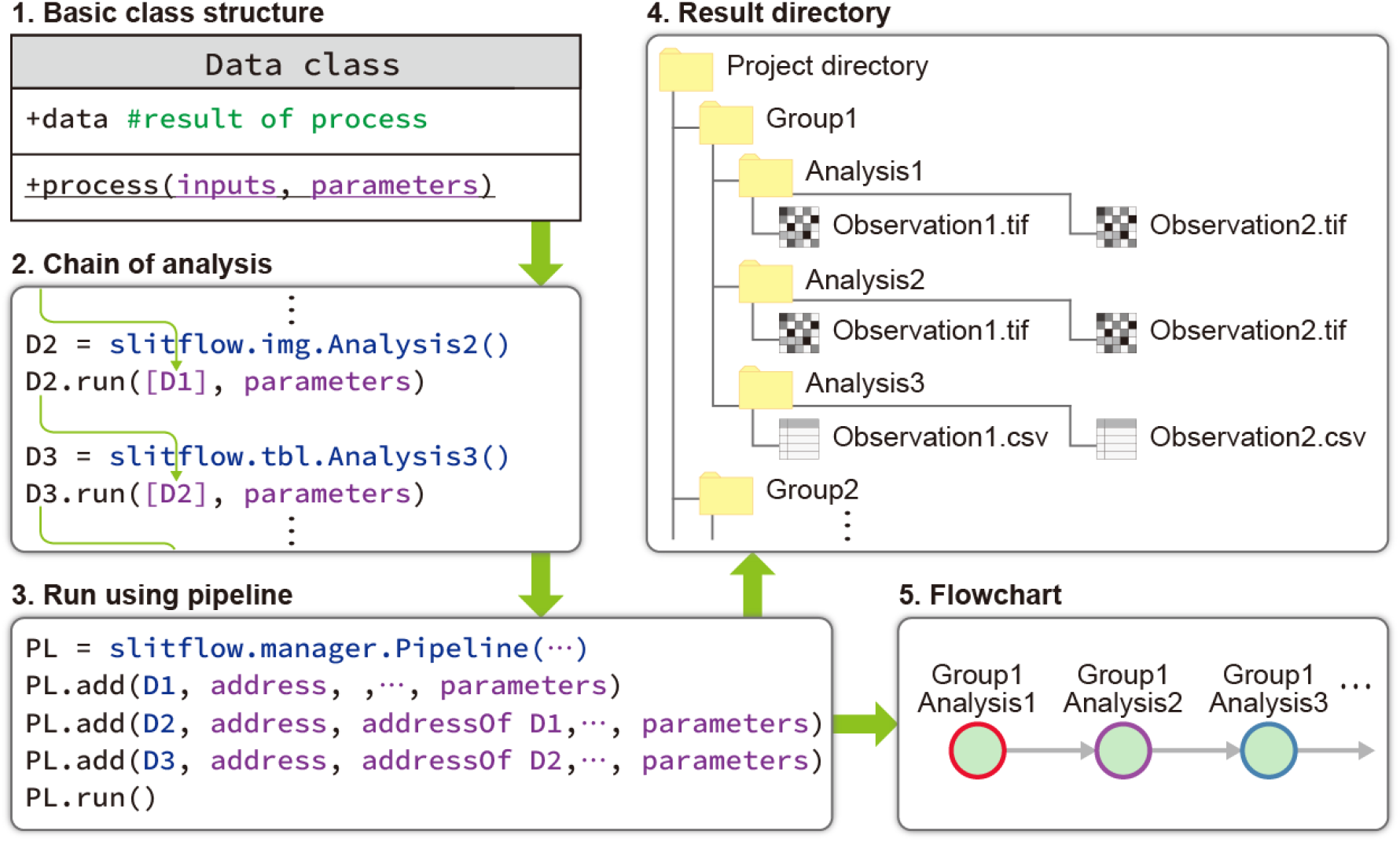
Simplified schematic of the Slitflow architecture.

#### 2.1.1. Reproducibility

An ideal reproducible workflow should obtain the same results without additional processing or parameter completion based on the reproducer’s intentions, independent of the computing environment. Slitflow focuses on four aspects that tend to result in low reproducibility in single-molecule analyses.

- *Task*: Slitflow requires the creation of a Data class for any trivial task. This includes selecting and sorting the data, performing simple arithmetic operations, and changing the line thickness of the resulting figures.
- *Parameter*: All parameters required for processing are entered each time as a dictionary of name and value sets. These parameters are exported along with the results and reused during reproduction. They are also readable by non-software users with accompanying parameter descriptions.
- *Pipeline:* All Data classes used in the workflow, their parameters, and dependencies of the resulting data are described as a pipeline script in the CSV file. Users can quickly reproduce, modify, and distribute the pipeline.
- *Environment:* Slitflow is written in Python, is freely available, and can be used in Windows, macOS, and Linux operating system environments. Users can run the analysis on computers with small memory capacities or large multithreaded computers by simply changing the loading and processing unit size to suit their computing environment.

#### 2.1.2. Accessibility

Inaccessible black-boxed analysis methods and intermediate results may prevent workflow validation and expansion. Slitflow aims to build an open workflow by designing access means for the following three elements:

- *Process:* Individual tasks can be run independently of the workflow and used as functions in user scripts to evaluate the functionality or implement it in their own algorithms.
- *Format:* Slitflow does not recommend creating proprietary data formats. An ideal data format should be opened and interpreted by users using standard applications initially installed on an operating system. If this is difficult, Slitflow recommends creating a Data class that converts the format into images, tables, or text for viewing.
- *Result:* The Slitflow pipeline maps all Data classes to the resulting folders. All tasks are based on loading the required data file and saving the resulting file in a subfolder. This system allows users to access and examine all the intermediate data of any task in the workflow.

### 2.2. Software functionalities

#### 2.2.1. Data

To create an analysis workflow using Slitflow, users must write all tasks as subclasses of the Data class, which is the basic unit of a data-processing pipeline. The Data class is categorized based on the resulting data type—such as the NumPy ndarray for image data and the Pandas DataFrame for table data. Slitflow provides various Data classes, including data loading, image processing, trajectory analysis, localization analysis, and figure creation. Users can find a Data class that fits their purpose from the API documentation. Users can also create a user-defined Data class by temporarily defining it in script code or creating a submodule file in the user subpackage. The Data class has the following structure to ensure scalability in data splitting, parallel processing, and workflow reuse:

#### 2.2.2. The process() method

The Data class has a static process() method that is directly accessible without creating a Data object. This structure enables users to execute any task independently of the workflow as follows:

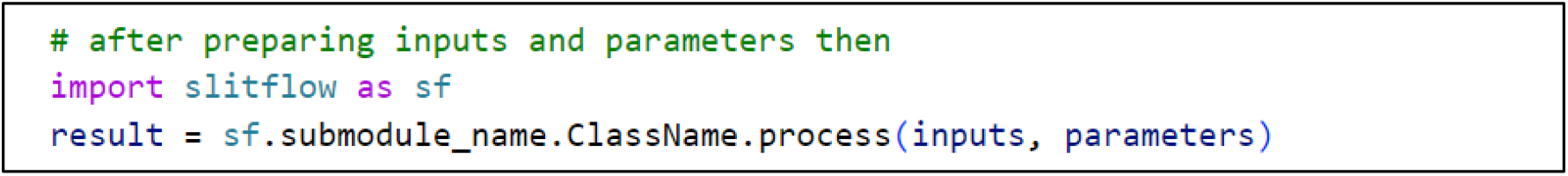

The process() method requires input data and a parameter dictionary. The input data comprises a list of actual data directly used in the analysis. Each list element corresponds to the result of a different Data object when the process() is called from the pipeline.

The following script creates the *x* and *y* coordinates of a two-dimensional random walk. This class requires an index table to add trajectories—such as images and trajectory numbers. This class also requires the diffusion coefficient, time interval, total trajectory length, and column names of the coordinates. The required input data and parameters can be found in the API reference in the software documentation.

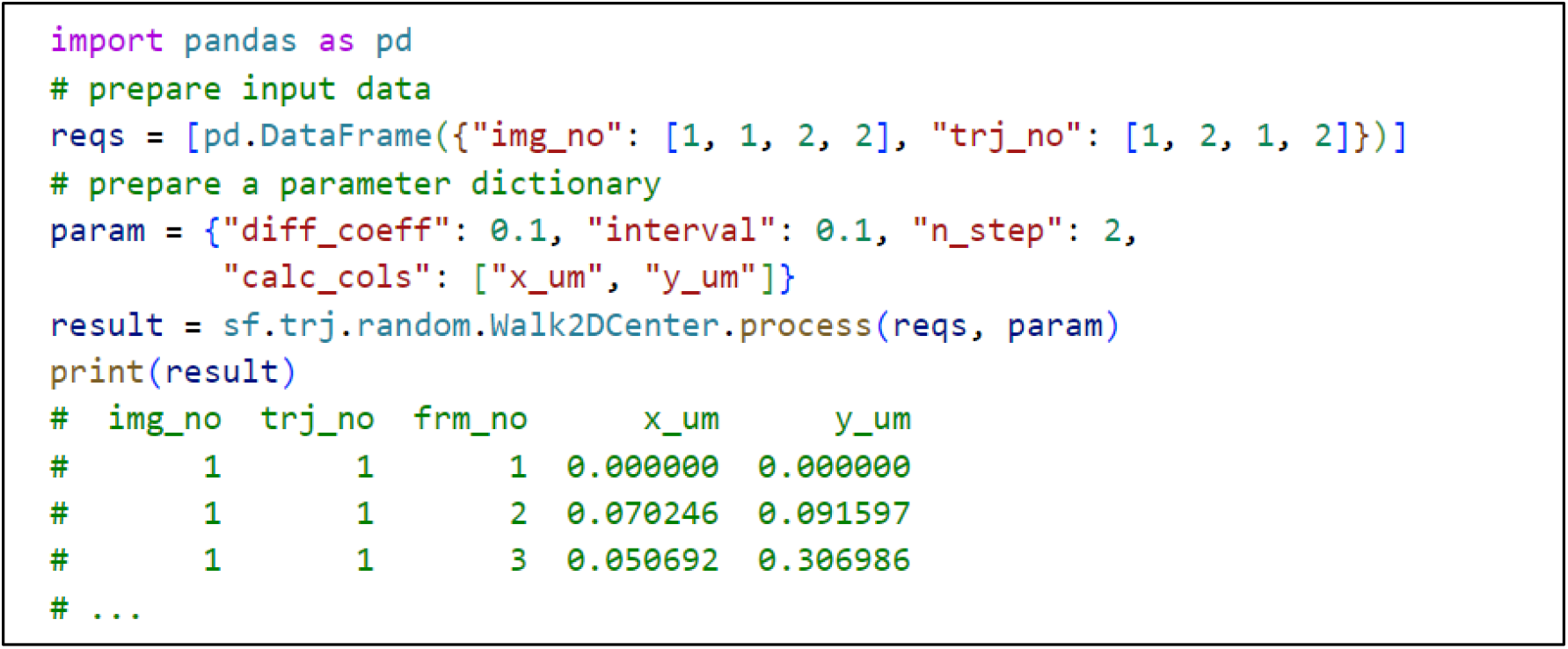

#### 2.2.3. The run() method

The static process() method is executed from the run() method if the users create a Data object for the analysis workflow. The resulting data are stored in the data property of the object. The run() method requires a list of Data objects as input, instead of the raw data required for the static process() method. The following code executes the random walk calculation as a Data object. The input DataFrame is replaced with an index data object, which creates a table of nested indices.

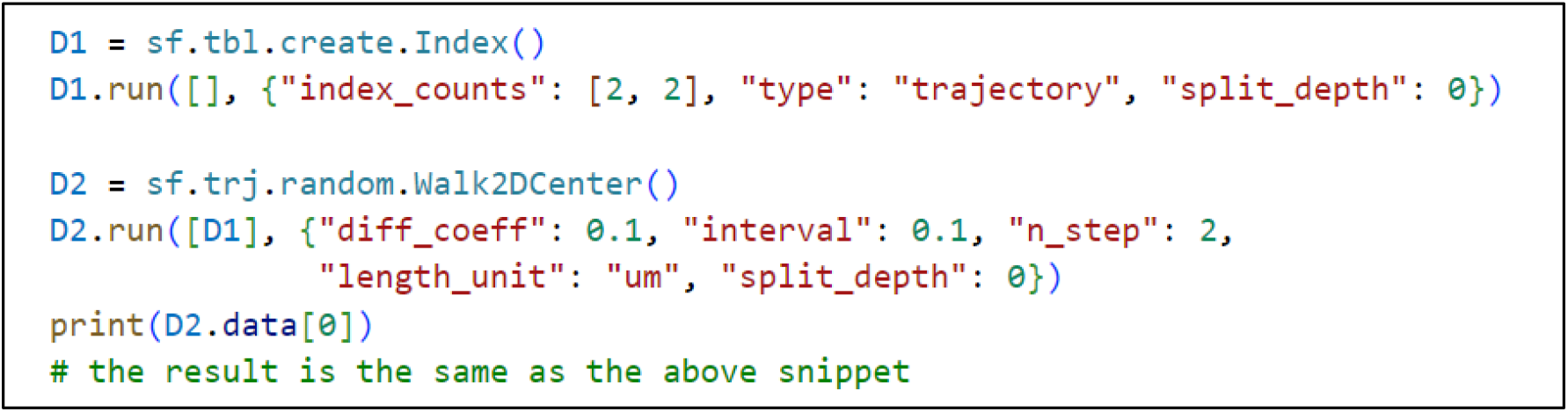

#### 2.2.4. Info and index

The run() method executes not only the process() but also other preparation steps— including adding parameter descriptions, creating a data index, and pairing different input data. Information about the parameters and the resulting data structure is stored in the info property of the Data object. This information is exported as a JSON text file that can be read by non-software users. The Data object also contains a table of data hierarchies— such as images and trajectory numbers. This table is used to pair different input data types and identify the selected images and trajectories. The following code snippet shows the information and index of the above random walk calculation Data object.

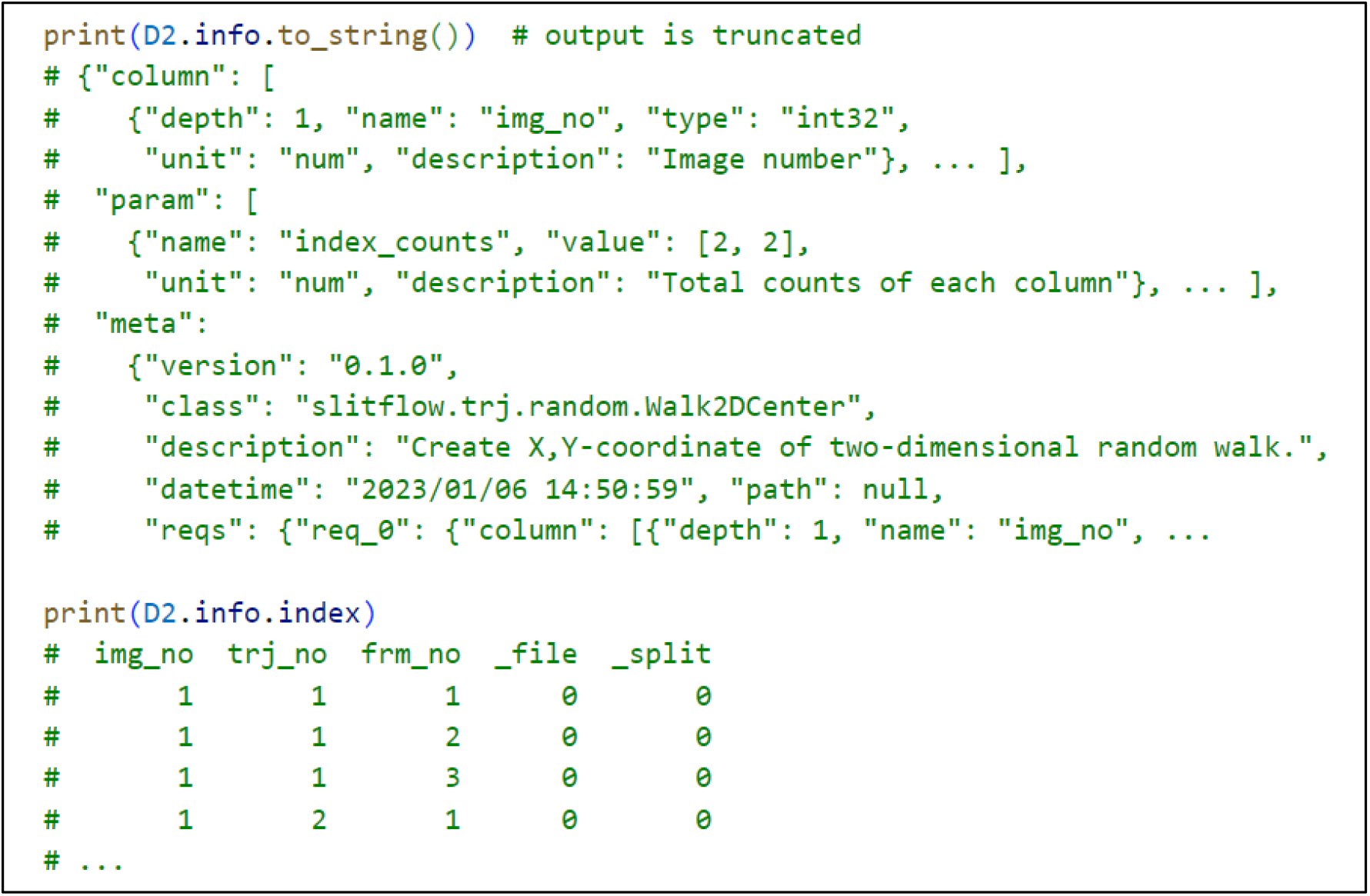

#### 2.2.5. Split depth

The data to be analyzed have a hierarchical structure—for example, the trajectory number, image number, replication number, and observation number. When applying analysis to data or dividing and saving data, it is necessary to specify the hierarchy in which the analysis or saving is to be performed. Slitflow allows users to change the target hierarchy of tasks easily by specifying the split depth. For example, users can calculate the average coordinates for each trajectory and output them in a single DataFrame. Using the same input, users can also average the trajectory coordinates for each image and export them to a DataFrame for each image by changing only the split depth. Additionally, users can use run_mp() instead of run() to compute the split data using multiple processes, which accelerates the calculation process by means of parallel computing.

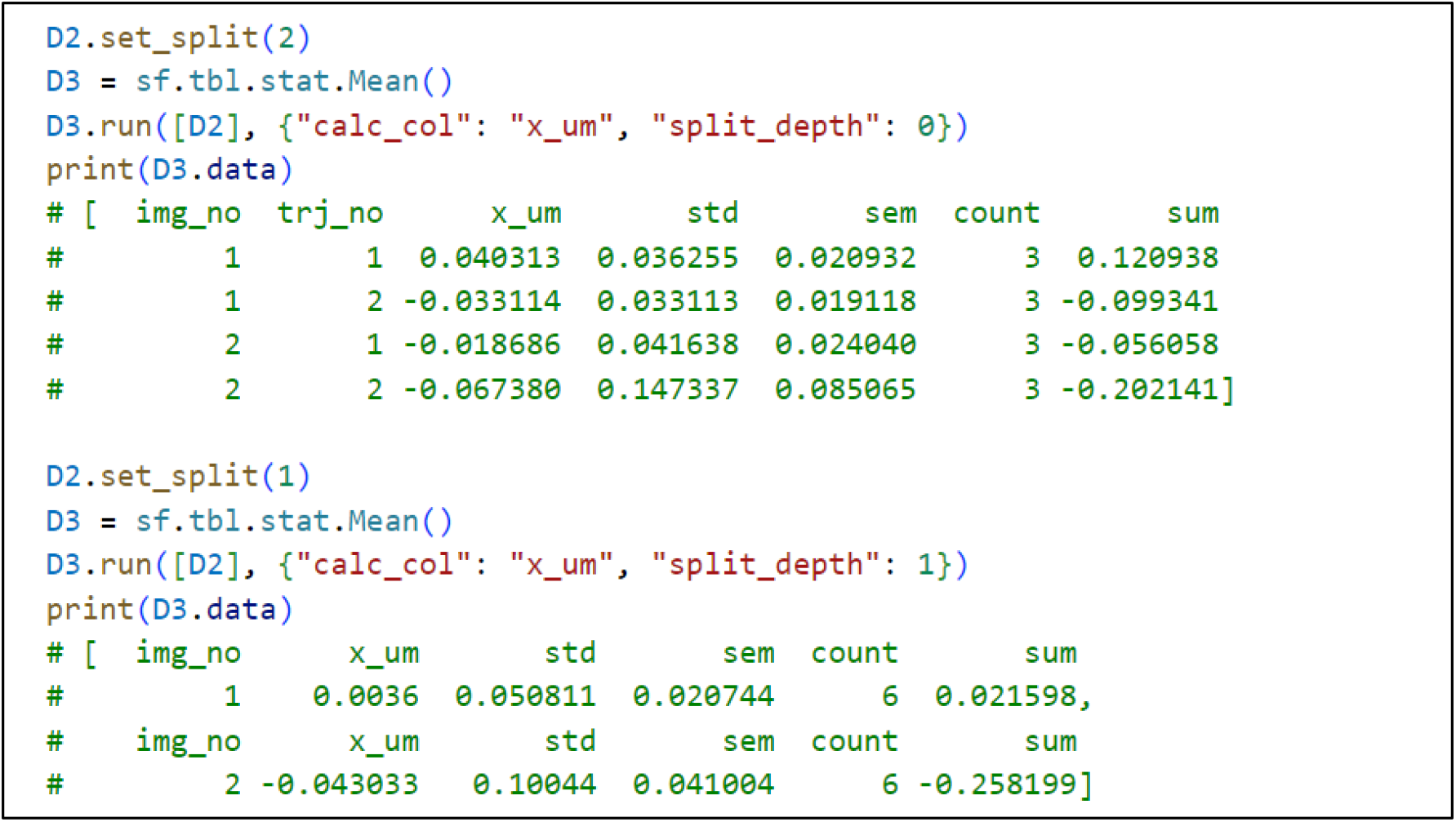

#### 2.2.6. Save and load

The resulting data are exported to the file format to which the Data class belongs by simply executing the save() method for the Data object. The data are divided into files according to a specified split depth. Indexes—such as the image and trajectory numbers of all split data—can be viewed in the text file exported with the resulting data, in which the data hierarchy is described.

#### 2.2.7. Pipeline

While the Python script and Jupyter notebook can be used to create an analysis pipeline script by connecting a series of Data objects, Slitflow provides a pipeline system for improving the workflow organization by standardizing the data-exporting directory. The Pipeline class creates a hierarchy in the project directory—comprising group folders and analysis subfolders—with each task corresponding to an analysis subfolder. Individual Data objects are stored in units of observation linked to file names throughout the subfolders (**Fig. 1**). Users select the folder location for the required and resulting data of the task by specifying an address comprising a group number and an analysis number. After inputting the additional parameters and observation names, the entire workflow is constructed by adding Data objects to the Pipeline object. All analyses are performed for all observations by executing the Pipeline run() method as described below.

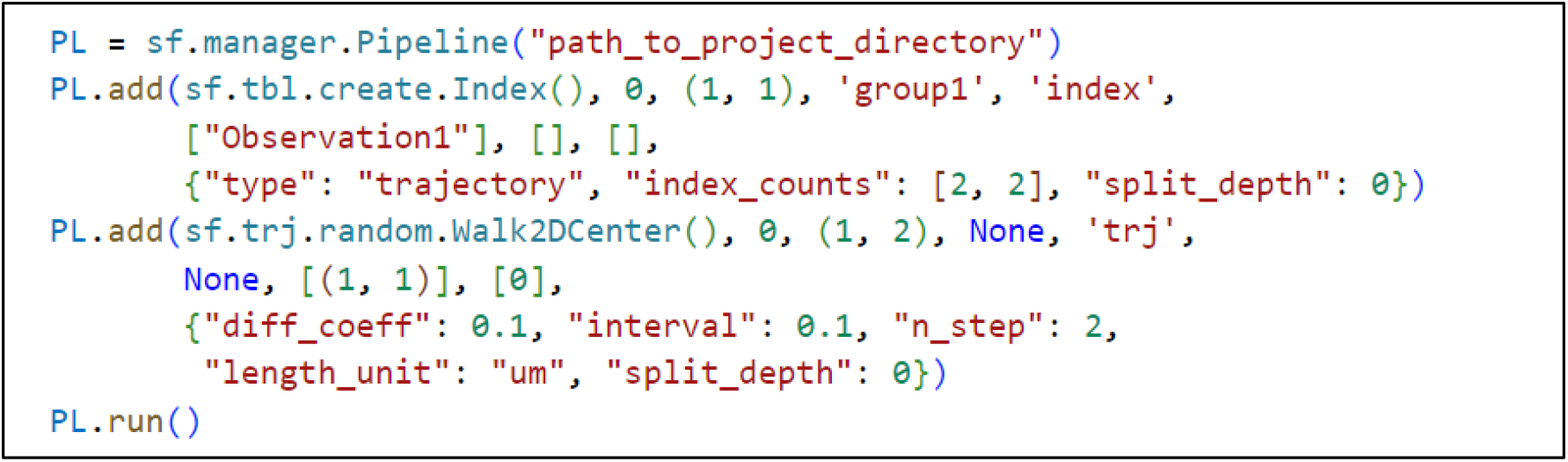

The created workflow can be output as a CSV file and reused for additional observations, rewritten parameters for adjustment, or distributed to other researchers. Researchers who receive the workflow file can execute a three-line script, as shown below, to fully reproduce a complex analytical workflow.

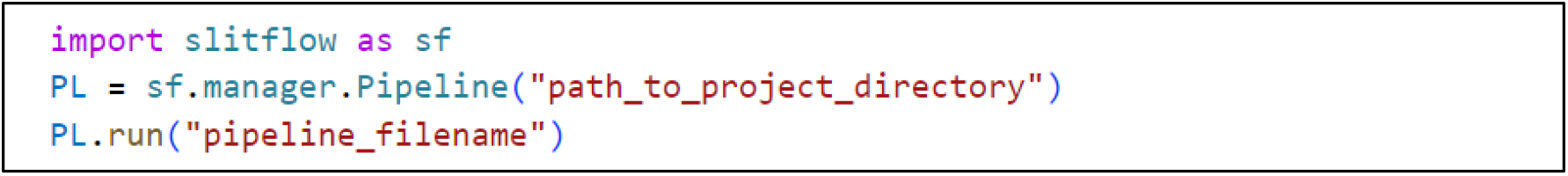

The data dependencies in the pipeline can also be used to create a flowchart, as shown in **Fig. 3**.

## 3. Illustrative examples

To demonstrate single-molecule analysis using Slitflow, we used live-cell single-molecule movies of RNA polymerase II (Pol II), a transcriptional protein complex, from cultured cells. Human U2OS cells expressing Halo-RPB1—the largest subunit of Pol II labeled with a self-labeling HaloTag—were stained with 3 nM Janelia Fluor 549 fluorescence substrate, and excited using HILO illumination [4]. 300 frames were captured at 33.33 ms per frame using an EMCCD camera. Single-molecule movies were used to extract and track bright spots and to visualize and analyze the trajectory on a single pipeline script.

### 3.1. Tracking

Single-molecule tracking requires pre-processing and tracking algorithms that are appropriate for the characteristics of the acquired images. In this study, we implemented a multistep customized process that focused on improving the location accuracy and processing time. First, fluorescent spots were detected using a Difference of Gaussian filter and the local maximum—as used in u-track [9] and TrackMate [8]—and then selected using a cell nucleus region mask and an intensity threshold. The positions were further refined by 2D Gaussian fitting using the *scipy*.*optimize*.*curve* fit, the trajectories being extracted using the *link* function of Trackpy [15]. To exclude noise trajectories, those with at least nine steps were selected. The detected trajectories were exported as tiling images using a series of drawing classes (**Fig. 2(a)**).

**Fig. 2.**
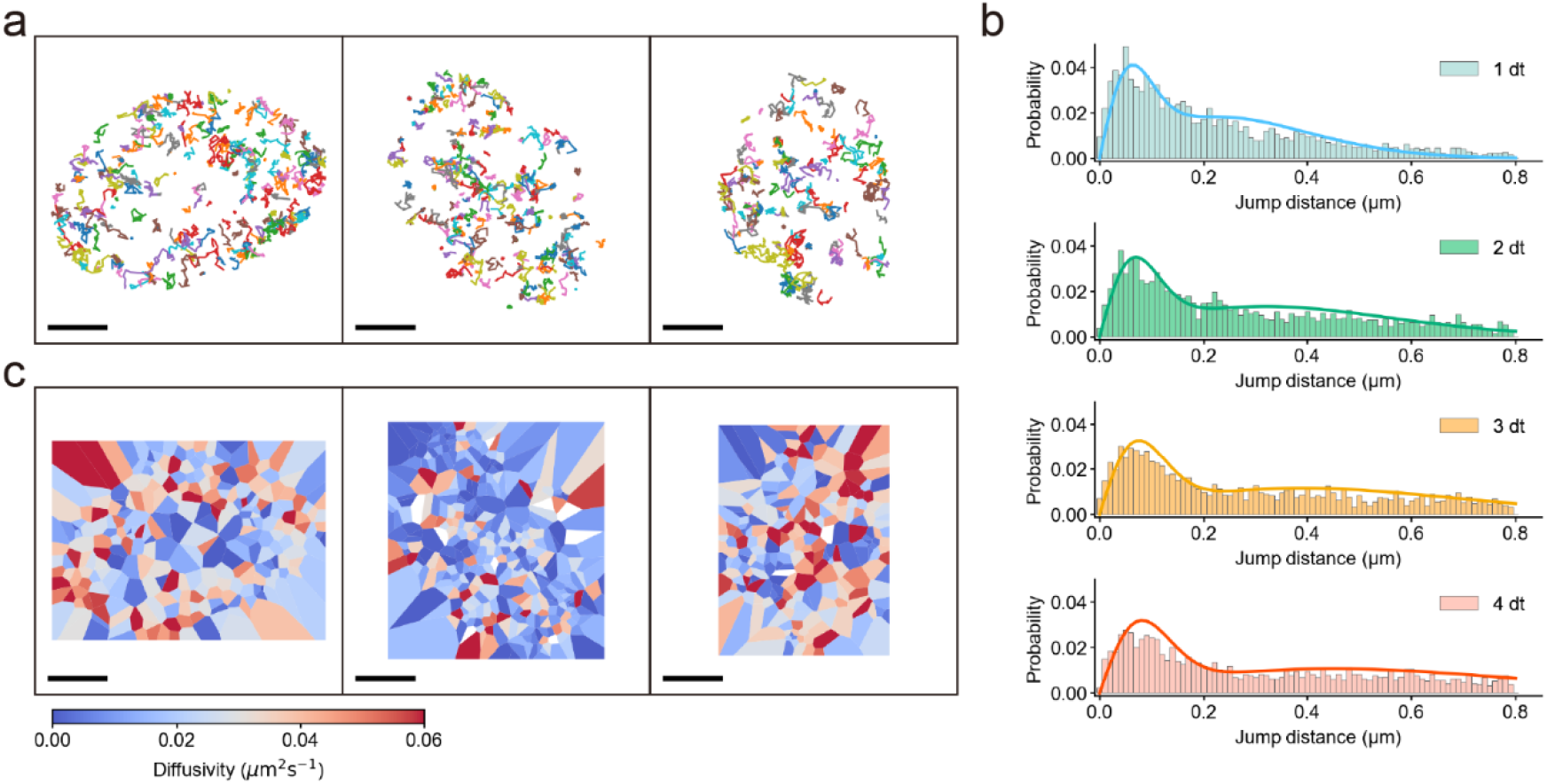
Example results from a single-molecule integrated workflow. The detected trajectory image (a), fitted jump distance distribution of Spot-On (b), and diffusivity map with TRamWAy (c) of Halo-RPB1 in U2OS cells are rendered. Bars, 5 μm.

**Fig. 3.**
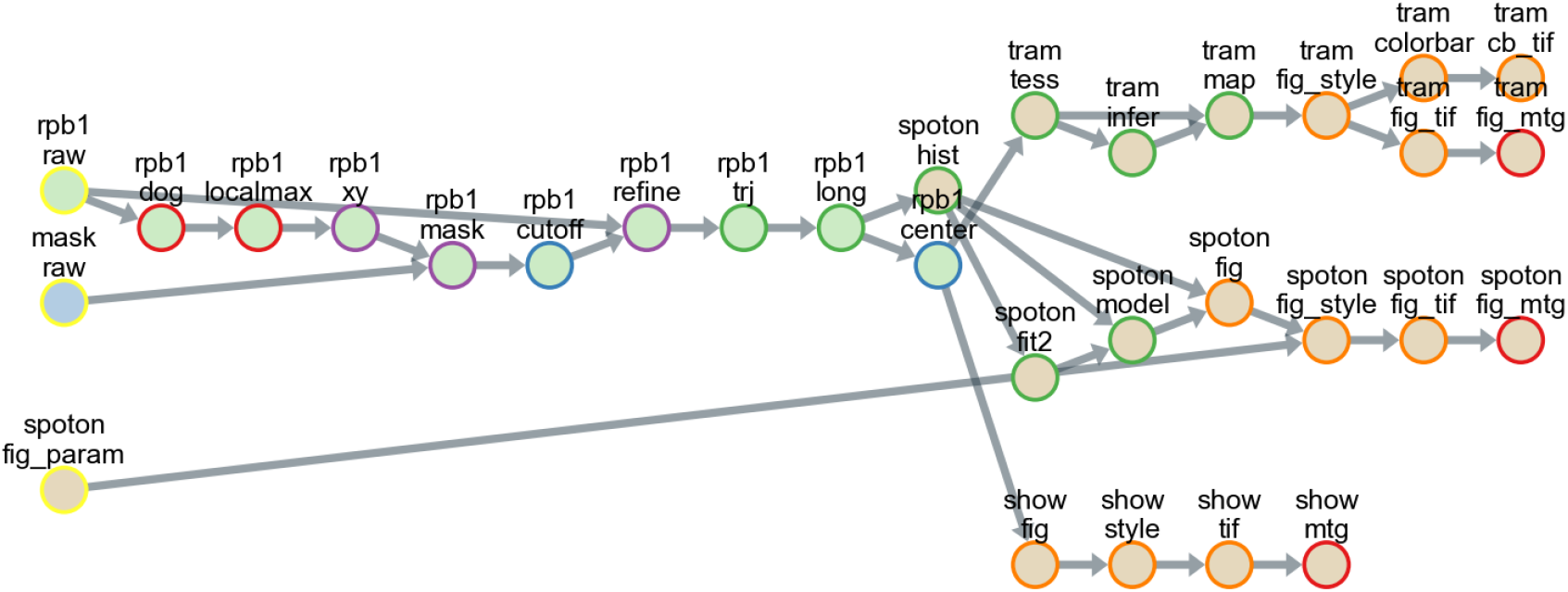
A flowchart of the entire single-molecule analysis, including tracking, state-of-the-art analyses, and figure creation.

### 3.2. Analysis

Methods for analyzing single-molecule trajectories are currently being developed, and many of them are available in the Python environment. In this example, two state-of-the-art analyses—that is, Spot-On [11] and TRamWAy [16]—were applied to detected Pol II trajectories. A simple wrapper class was implemented for each analysis with the data format and figure output adjusted to match the Slitflow architecture. Using these classes, the jump-distance histograms and their fitting parameters for the dynamic component were calculated using Spot-On (**Fig. 2(b)**). Furthermore, a spatial map of the diffusivity of Pol II molecules was obtained from the trajectory localization information using TRamWAy (**Fig. 2(c)**).

### 3.3. Flowchart

All tasks, including tracking, analysis, and drawing, were saved as a single pipeline script text file in the CSV format for reuse and distribution. Using the pipeline script, a series of data-processing steps from the raw data to the final image could be exported as a flowchart (**Fig. 3**), each circle in the flowchart representing an individual task corresponding to an analysis subfolder in the project directory. The arrows between circles represent data dependencies. In this example, 26 different classes were used, and all the data were stored in 31 subfolders in five groups.

## 4. Impact

Slitflow systematically provides reproducibility and accessibility for the analysis of diverse and complex data and methods, enabling robust integrated analysis. Integrated single-molecule analysis, as demonstrated here, greatly benefits the molecular and cellular biology research community. The comprehensive concept underpinning the Slitflow architecture can also serve as a basic guideline for constructing workflows in many research fields that deal with scientific data. The flexible data structure is applicable not only to single-molecule imaging, but also to image analysis, numerical simulation, and many other research fields in which data arrays and tables are repeatedly processed. We demonstrated customized tracking by combining u-track [9] and Trackpy [15] logic with state-of-the-art analyses of trajectory [11] and localization [16] in a single workflow. This integrated functionality allows tracking and localization verification—as reported in competitions [17, 18]—to be performed locally in a single workflow. Additionally, integrated analyses combined with cutting-edge studies can be immediately distributed to the scientific community using the pipeline script. Furthermore, the accumulation of user-defined classes paves the way for constructing cross-field analysis methods—such as spatial genomics [19]—and applying deep learning to various processing steps [20, 21] for single-molecule imaging.

In practice, developers can concentrate on creating methods without considering file management, pre-processing, and visualization. Developers can easily create various types of figures in publication quality. Owing to the readable pipeline scripts and class documentation, users with no programming experience can understand and execute individual processes. These features facilitate productive communication between developers and users as well as analysts and collaborators. Moreover, Slitflow has implemented many single-molecule analysis methods that our group has published with collaborators [22, 23, 24]. We shared the software and its results with several research groups and continue to develop it. The latest software can be installed using the Python Package Index with its extensive documentation and examples. The public GitHub repository allows anyone to contribute to debugging and adding functionality, and we are confident that this software will be widely used by the research community.

## 5. Conclusions

Slitflow is a flexible Python framework that provides a complete workflow for complex single-molecule analyses using a comprehensive architecture based on reproducibility and accessibility. This package creates a series of processes, including image pre-processing, tracking, localization analysis, data sorting, and figure creation, by connecting simple task objects that correspond to each result file. Slitflow was designed by considering users who developed analysis tools, validated multiple analysis methods, reproduced workflows without programming skills, and used the results without installing software. The reusability and scalability of Slitflow could promote the development and deployment of new methods in various research fields that process scientific data.

## Acknowledgments

This work was supported by the Ministry of Education, Culture, Sports, Science and Technology/Japan Society for the Promotion of Science, Kakenhi (JP18H05527 and JP20K15755 to Y. Ito; JP19H03192 and JP22H02581 to M. Tokunaga).

## Data availability

Online documentation is available at https://slitflow.readthedoc.io/en/latest/. The data used in this study are available at https://zenodo.org/record/7645485#.Y_w1QR_P2Hu.

## Supplementary Materials

**Appendix**.

Supplementary material includes detailed descriptions of the run mode and the user-defined class.

### Run mode

The run mode is a Pipeline argument that specifies split-file loading and parallel computing. When adding a Data object to the Pipeline, the run mode is set to a value between zero and three. Modes 0 and 1 read all split files into memory simultaneously before executing the processing. Conversely, Modes 2 and 3 repeat the loading, processing, and saving cycle for each split data. Additionally, Modes 0 and 2 compute the data list sequentially using a single process. Conversely, Modes 1 and 3 compute the elements of the data list in parallel using different processes. **Table 1** shows the features of the run modes and tasks suitable for each mode.

**Table 1.**
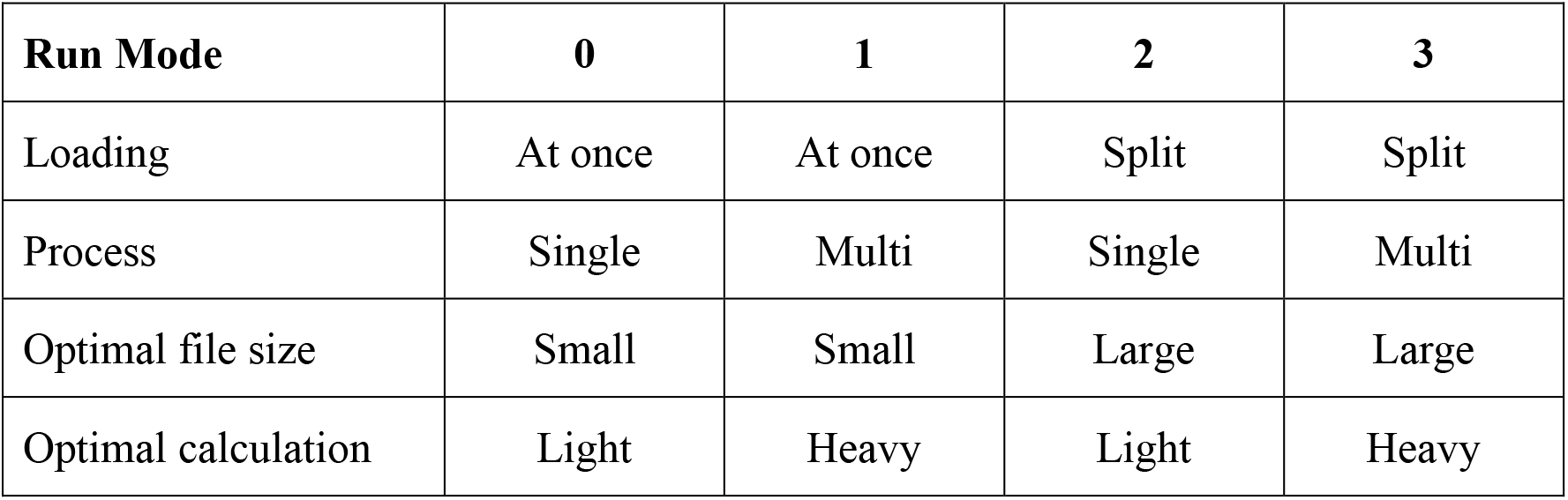
Features of the run modes and tasks suitable for each mode.

| Run Mode | 0 | 1 | 2 | 3 |
| --- | --- | --- | --- | --- |
| Loading | At once | At once | Split | Split |
| Process | Single | Multi | Single | Multi |
| Optimal file size | Small | Small | Large | Large |
| Optimal calculation | Light | Heavy | Light | Heavy |

For example, Mode 0 is used for the light processing of data or aggregation of all data. Mode 1 is used when the entire split data can be loaded into memory simultaneously; however, individual processing is too heavy, and parallel computation is preferred for each element of the data list. Mode 2 is used when the overall data are too large to load, but there is a computational overhead in processing the data list, or when parallel computation is used to process single split data. Mode 3 is used when big data are split and loaded and further parallel computation is required.

### User-defined class

A new task can easily be added to Slitflow as a user-defined class. The first approach is to define a class directly in the Python script or Jupyter notebook inherited from a Slitflow Data class. The script below defines a simple class in the executable script that adds one to the values of the table column, a class object being directly connected to a series of Data objects. This procedure does not enable user-defined classes to be added to the pipeline and run; however, it helps with new prototype classes.

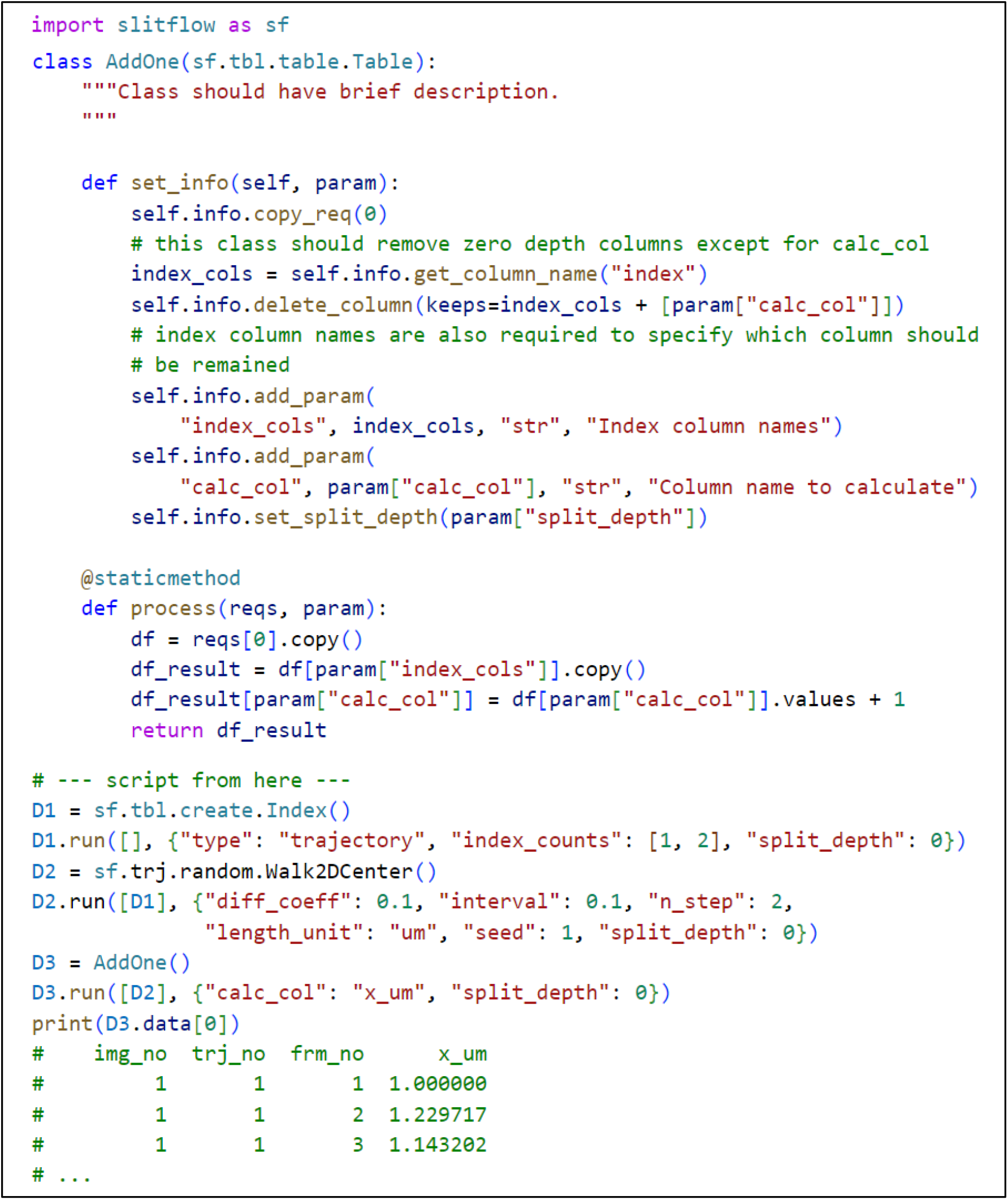

The second approach is to add a class as a module file to the user subpackage folder of the Slitflow source code. This allows the user-defined class to be added to the pipeline and incorporated within a reusable analysis workflow, or to create a flowchart. The following example executes the AddOne class of a template module saved as an example in a user subpackage.

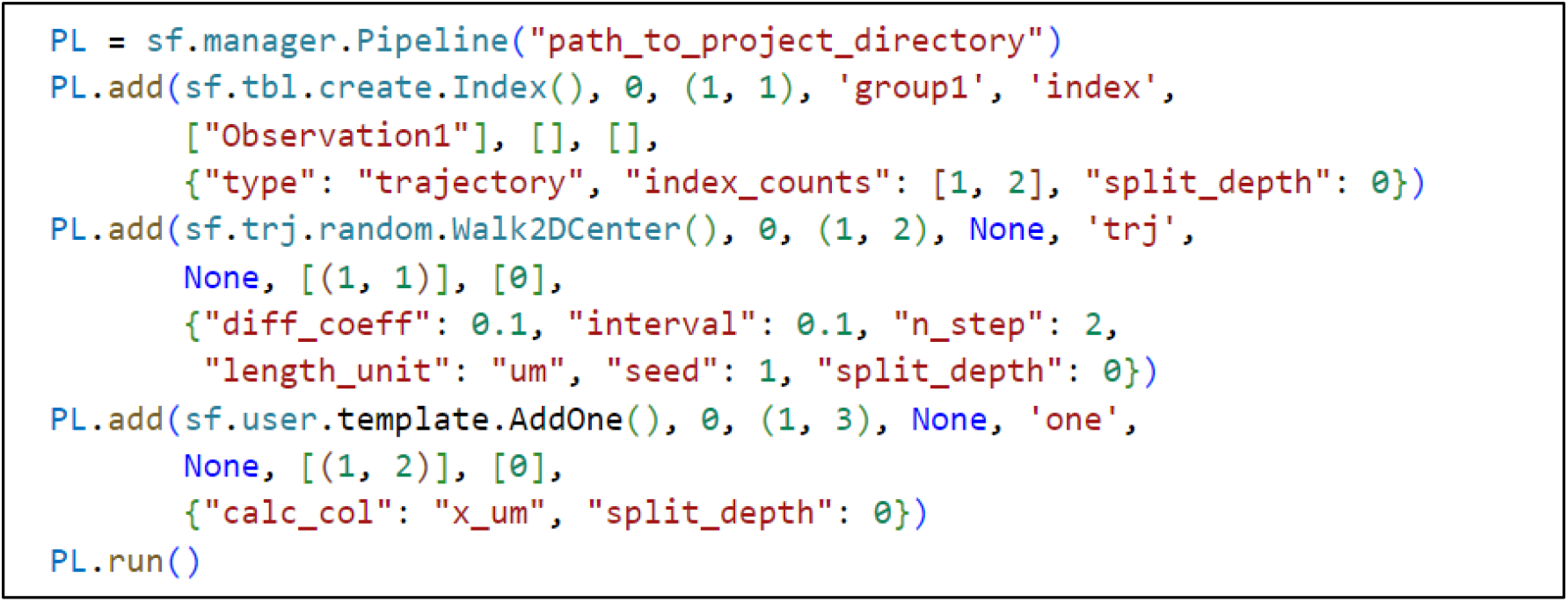

For more useful user-defined module files, consider moving them to the dev subpackage folder and pulling requests to the develop branch of the Slitflow repository to share them among users.

